## Supplemental Figures for "Ratios in Disguise, Truths Arise: Glycomics Meets Compositional Data Analysis"

[*]Corresponding author

### Supplementary Figures

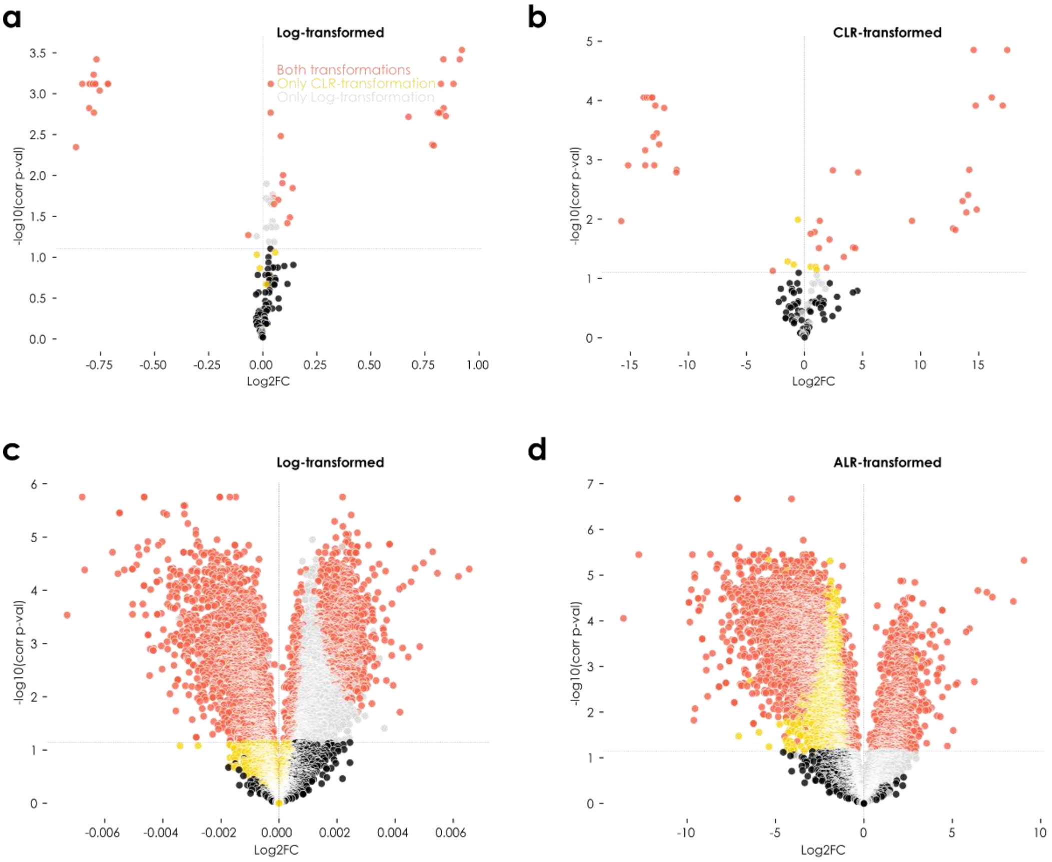


**Supplementary Figure 1. Analyzing glycoproteomics data via CLR-transformation. a-d)** Immunoglobulin glycoproteomics data from human milk (a-b, colostrum vs mature, Wang et al., *J Agric Food Chem*, 2021) or two liver cancer cell lines (c-d, Hep3B vs 97L, Kong et al., *Nat Commun*, 2022) were analyzed with the *get_differential_expression* function in glycowork (version 1.3), with either $log_2$-transformed relative abundances (a, c) or ALR/CLR-transformed relative abundances (b, d). Shown are volcano plots, where glycopeptides are colored red if both transformations identified their differential expression and yellow if only CLR-transformations recovered their differential expression. No glycopeptides were identified by log-transformation but not by CLR-transformation. The horizontal line indicates the sample-size appropriate significance thresholds of alpha = 0.08 or 0.07, respectively.

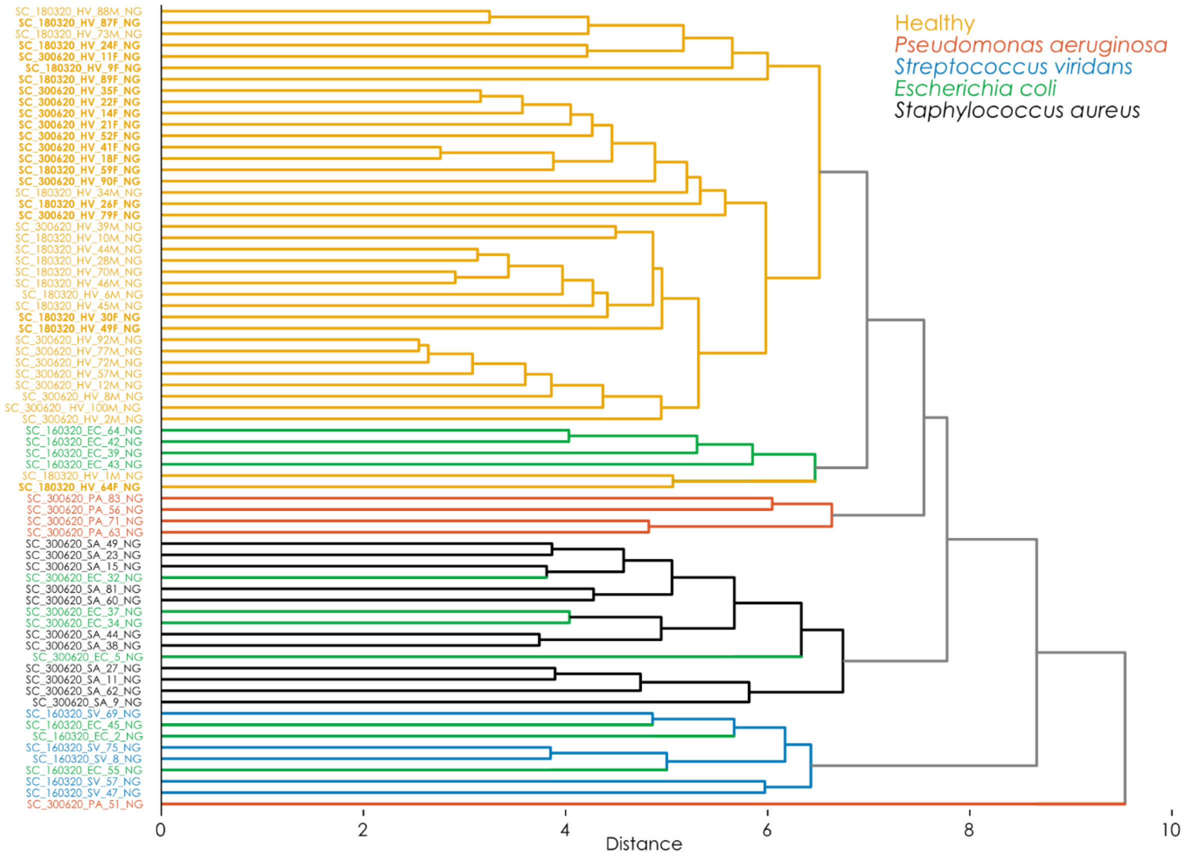


**Supplementary Figure 2. ALR-transformed data yield superior clustering.** Serum *N*-glycome data from healthy volunteers or bacteria-infected patients (Chatterjee et al., *J Clin Med*, 2021) were ALR-transformed via the *get_heatmap* function (glycowork, version 1.3). Then, an unsupervised hierarchical clustering was performed via the UPGMA algorithm on Euclidean distances. Classes are indicated by their color and, for healthy volunteers, female participants were indicated by bolding their ID, demonstrating even a clustering within the class by the known glyco-modifier sex.

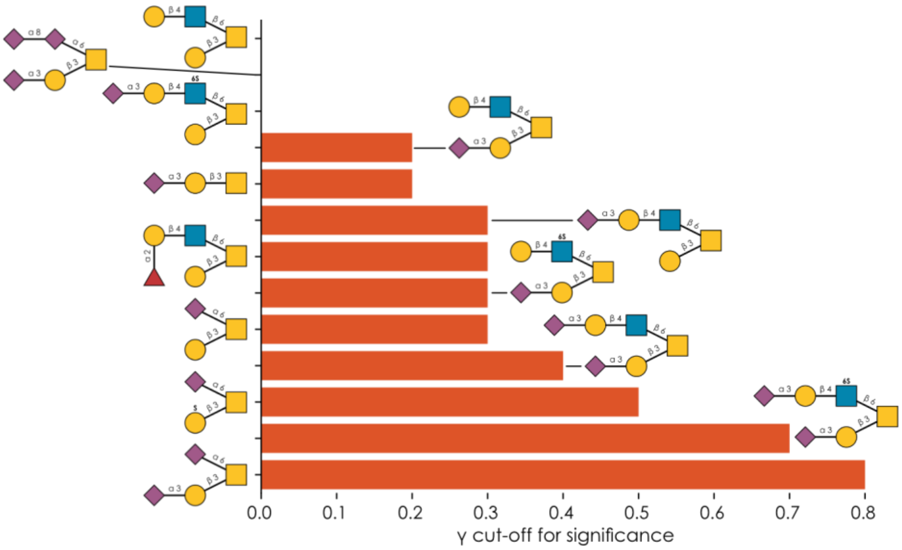


**Supplementary Figure 3. Scale uncertainty as a measure for effect robustness.** For the Basal Cell Carcinoma *O*-glycomics dataset (Möginger et al., *Front Oncol*, 2018), we calculated the differential expression between paired healthy and cancerous tissue (n = 40) using the *get_differential_expression* function (glycowork, version 1.3) and different values of γ as scale uncertainty, ranging from 0 to 1. Recorded as a bar graph, for each structure, is the value of γ at which no significant difference could be established (p < 0.044). Greater values of this γ-cutoff then indicate more robust effects, as they persist even under great scale uncertainty. Glycans are depicted via the Symbol Nomenclature for Glycans (SNFG).

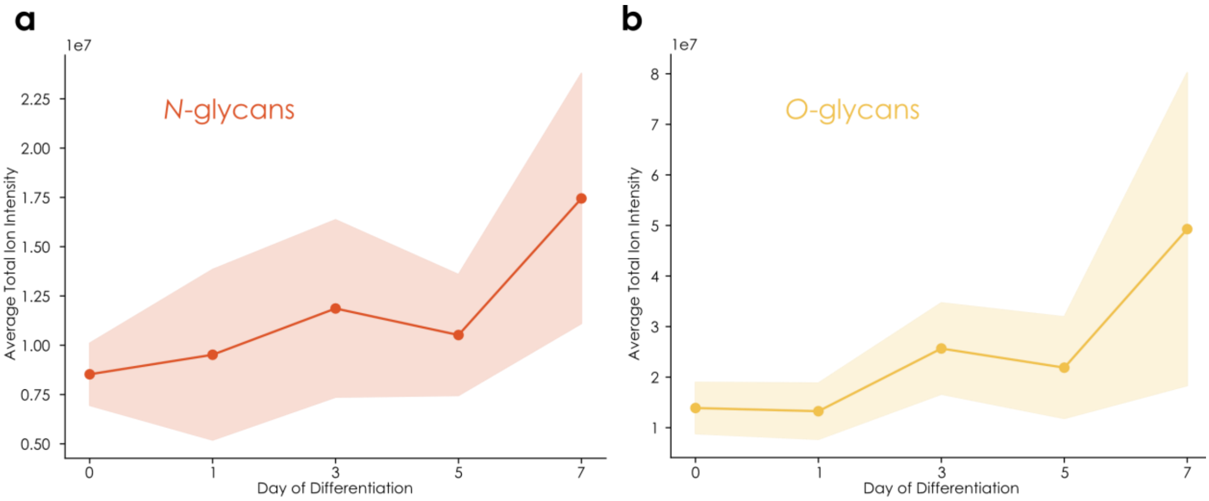


**Supplementary Figure 4. Change in total glycan signal over macrophage differentiation. a-b)** For each day of macrophage differentiation (Hinneburg et al., *Glycobiology*, 2020), for both *N*-glycans (a) and *O*-glycans (b), the integrated ion intensity of all glycans was summed for each sample and averaged across donors and replicates. Shown are the mean total ion intensity as a line plot, together with a 95% confidence band.

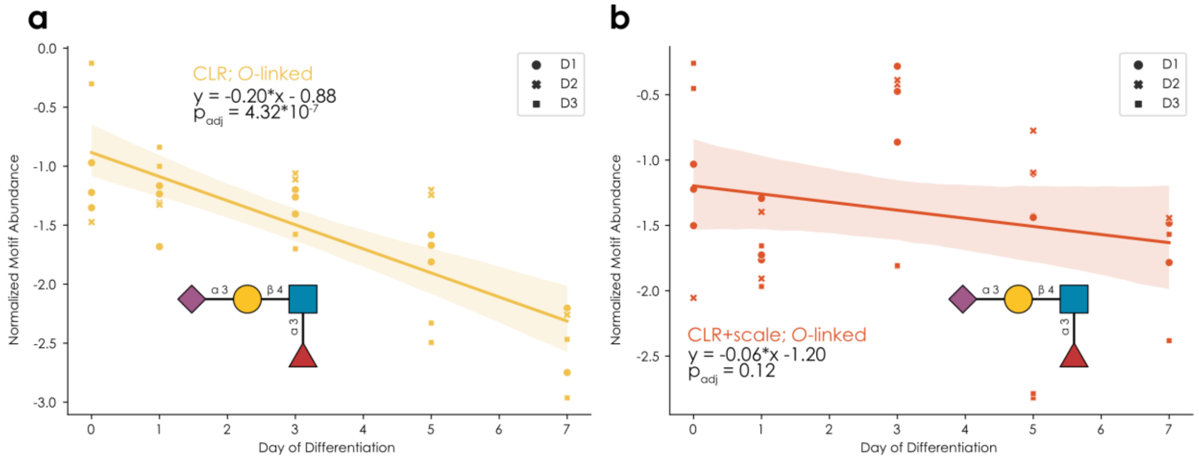


**Supplementary Figure 5. Relative and absolute decrease of sialyl-Lewis X in *O*-glycans during macrophage differentiation. a-b)** CLR-transformed O-glycomics data from a longitudinal macrophage differentiation dataset (Hinneburg et al., *Glycobiology*, 2020) was analyzed similar to Figure 3b-g, analyzing the temporal behavior of sialyl-Lewis X expression with a scale uncertainty model (a) or an informed scale model (b).

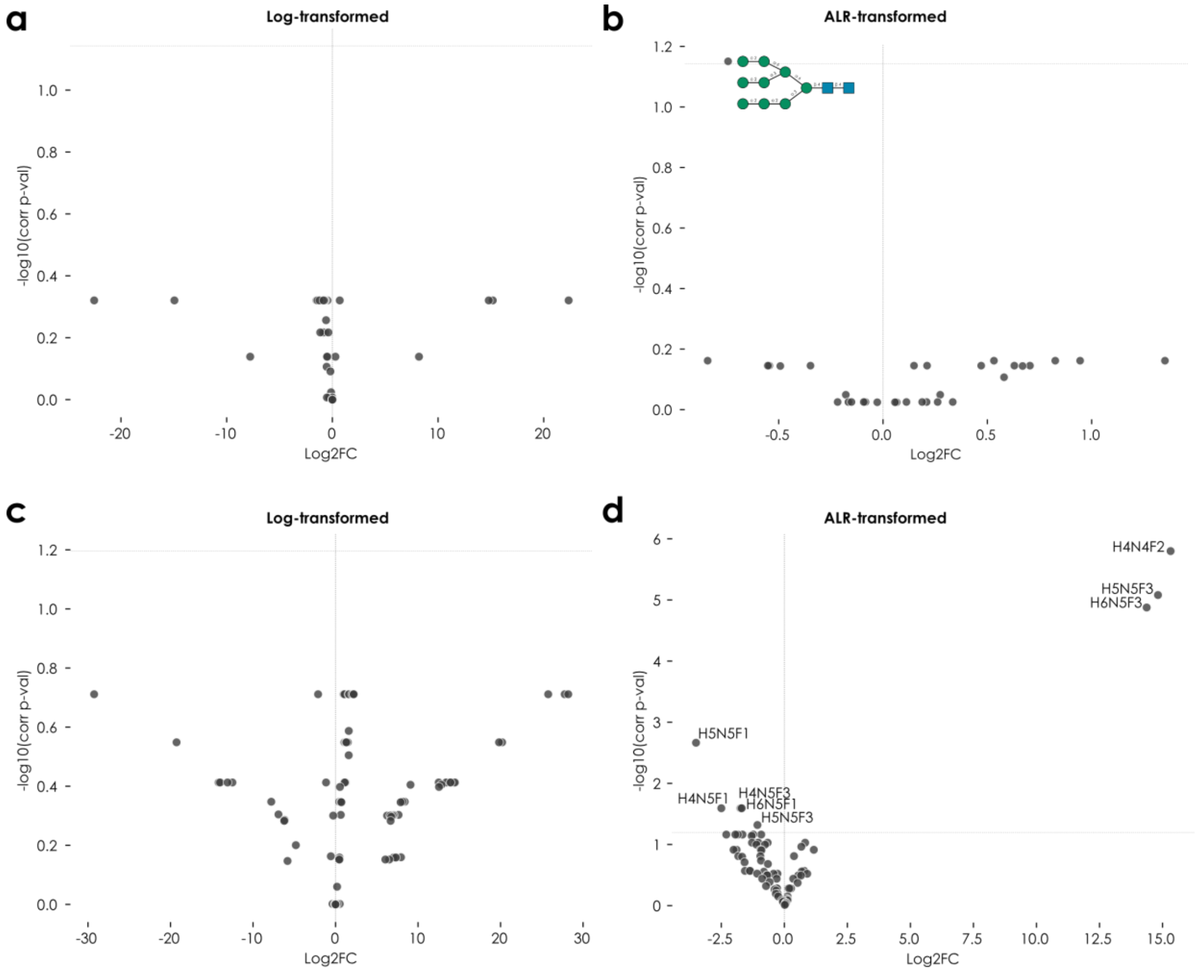


**Supplementary Figure 6. Directly using glycomics ion intensities for analyses suffers from low sensitivity. a-d)** *N*-glycomics data from a chronic lymphocytic leukemia (a-b; n = 8) and a colorectal carcinoma dataset (c-d; n = 12) from Chatterjee et al., *Oncotarget*, 2021, were used for differential expression analyses, using either log-transformed ion intensities (a, c) or ALR-transformed relative abundances (b, d). Shown are volcano plots, where a horizontal line indicates the alpha-cutoff for significant differential expression. Differentially expressed glycans are annotated with their SNFG representation or their composition if no structure was available.

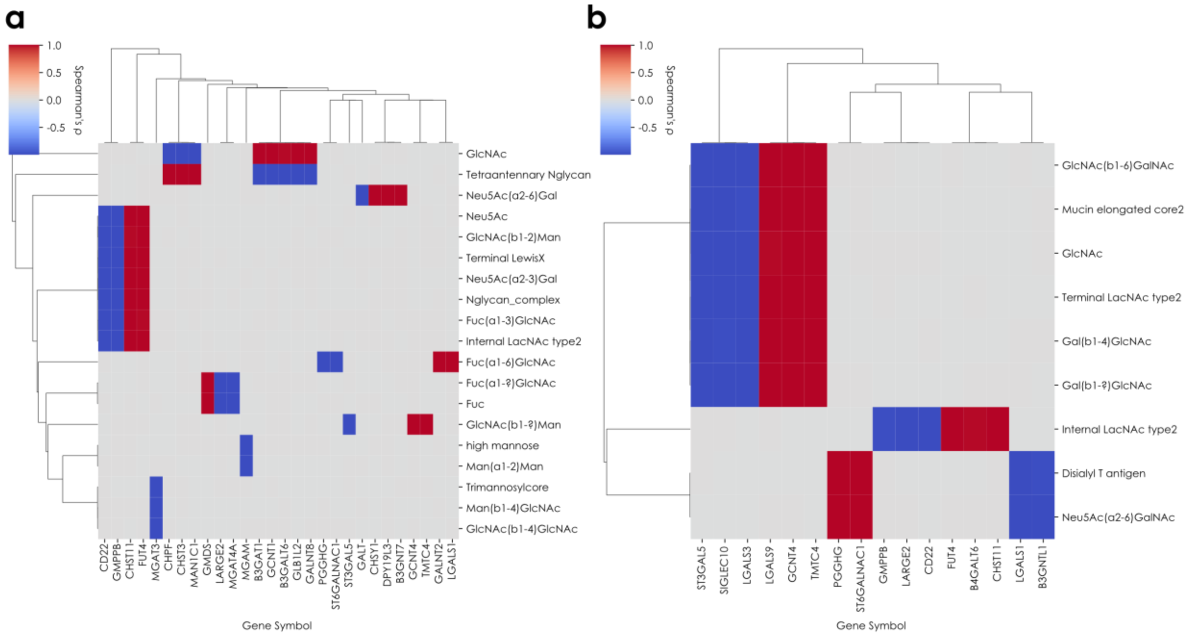


**Supplementary Figure 7. Cross-correlation between glycan motifs and transcriptomics using compositional data analysis. a-b)** Using B-cell glycomics and transcriptomics data from Oliveira et al., *Theranostics*, 2021, we filtered the list of differentially expressed for glycogenes, transformed both glycomics and transcriptomics data with CLR with a scale uncertainty model and calculated the cross-correlations between transcripts and *N*- (a) or *O*-glycan motifs (b). Shown are only significant correlations ($p < 0.083$, at a sample size of five), with their Spearman's rho as retrieved from the *get_SparCC* function (glycowork, version 1.3).
